# cRacle: R Tools for Estimating Climate from Vegetation

**DOI:** 10.1101/641183

**Authors:** Robert S. Harbert, Alex A. Baryiames

## Abstract

**Premise of the study:** The Climate Reconstruction Analysis using Coexistence Likelihood Estimation (CRACLE) method for the estimation of climate from vegetation is a robust set of modeling tools for estimating climate and paleoclimate that makes use of large repositories of biodiversity data and open-access R software.

**Methods:** Here we implement a new R library for the estimation of climate from vegetation.The ‘cRacle’ library implements functions for data access, aggregation, and models to estimate climate from plant community composition. ‘cRacle’ is modular and features many best-practice features.

**Results:** Performance tests using modern vegetation survey data from North and South America shows that CRACLE outperforms alternative methods. CRACLE estimates of mean annual temperature (MAT) are usually within 1°C of the actual when optimal model parameters are used. Generalized Boosted Regression (GBR) model correction is also shown here to improve on CRACLE models by reducing bias.

**Discussion:** CRACLE provides accurate estimates of climate from modern plant communities. Non-parametric CRACLE modeling coupled to GBR model correction produces the most accurate results to date. The ‘cRacle’ R library streamlines the estimation of climate from plant community data, and will make this modeling more accessible to a wider range of users.

## INTRODUCTION

‘cRacle’ is an open-source R software package that performs the Climate Reconstruction Analysis using Coexistence Likelihood Estimation (CRACLE) model for the estimation of climate from biological community data (Harbert & Nixon, 2015). CRACLE performs an empirical likelihood estimation of climatic parameters given a set of taxa (species, genera, or families) and their modern distributions via point-locality data from the Global Biodiversity Information Facility (GBIF) or similar primary biodiversity databases (e.g., iNaturalist, iDigBio, BIEN).

The primary application of CRACLE and related methods is in the reconstruction of paleoclimate from fossil floras. All current applications of CRACLE make assumptions about climatic niche stability through time because the rely on the Nearest Living Relative (NLR) approach to infer elements of fossil taxon niche dimensions by observing the niche space occupied by a closely related extant relative (Mosbrugger and Utescher, 1997; Utescher, 2014; Harris *et al*. 2014). The NLR assumptions will be most valid when the fossil taxa can be placed as members of extant species and least accurate when fossils are members of extinct lineages. For this reason, NLR and CRACLE will produce the most reliable reconstructions of relatively recent (e.g., Pleistocene and Pliocene) paleoclimate, but may produce reasonable results for much older fossil assemblages where the plant fossil taxa can be placed reliably in extant groups with well sampled modern distributions (e.g., Harris *et al.* 2014).

CRACLE modeling was recently applied to the estimation of paleoclimate using Late Quaternary (<50ka – *kilo anuum*) plant macrofossils from packrat (*Neotoma* spp.) middens in Western North America (Harbert, Nixon 2018). Analysis of the packrat midden paleoclimate estimates revealed a history of rapid climate change during and just after the Late Pleistocene deglaciation, followed by Holocene warm and dry periods of 1-2°C above the modern (1950-2000) averages (Harbert and Nixon, 2018)

Community assemblages from excavated permafrost at five Pliocene sites in the Canadian Arctic Archipelago analyzed with CRACLE (Fletcher *et al*., 2017). This work revealed that the Canadian arctic was up to 22°C warmer than today during the Early Pliocene (~3.6Ma – *mega anuum*) and supported many cool-temperate plant taxa. However, Fletcher *et al.* (2017) carefully compared results derived from CRACLE estimates using species only identifications vs. generic level identifications and suggest that care must be taken when interpreting CRACLE results when generic level identifications are used to avoid being misled by model imprecision.

Modified bioclimatic envelope method that are closely related to CRACLE were recently used to reconstruct the Paleocene-Eocene Thermal Maximum (~55Ma) and the Eocene Thermal Maximum (53.5Ma) by way of plant macrofossils from the Green River Basin in southwestern Wyoming and plant microfossils found in Arctic marine sediment cores (Hyland et al., 2018; Willard et al., 2019).

CRACLE models have been applied to non-plant communities as well. Pliocene Arctic climate estimates based on fossil beetle communities using CRACLE, revealed notable biases in beetle based climate reconstructions relative to plant proxy and stable isotope methods (Fletcher *et al*., 2018).

Despite CRACLE existing for the first four years as only a single R script published as a supplement to the original paper (Harbert and Nixon, 2015) the method has proven to be a useful tool to the community (presently 16 Google Scholar citations), yielding informative results for several fossil systems. Here, we present the ‘cRacle’ R package. ‘cRacle’ is designed to streamline the process of CRACLE modeling, data acquisition and pre-processing of primary biodiversity data (from GBIF and other related databases), and CRACLE result visualization. The ‘cRacle’ R package is built from modular functions that perform segments of the modeling protocol that allow for easy to read R scripts as well as future modular developments of related methods.

For comparison with the CRACLE methods, we have also implemented the related Thompson’s Mutual Climatic Range (MCR; Thompson et al., 2012). We include both the weighted and unweighted MCR methods as representatives of the many available MCR or Coexistence Approach (CA) range-intersect methods (Mosbrugger and Utescher, 1997; Sinka and Atkinson, 1999; Utescher et al., 2014) for the estimation of climate from vegetation (ECV) as part of the ‘cRacle’ R package. The ‘cRacle’ package is distributed under the open MIT License and all code is available on GitHub (https://www.github.com/rsh249/cRacle).

## METHODS

Full documentation of ‘cRacle’ R code is available with the package code (https://github.com/rsh249/cRacle). Below is an outline of the major functionality of the ‘cRacle’ package and explanation of many of the core functions available.

### ‘cRacle’ Package Functions – Data Acquisition

Downloading primary biodiversity data from GBIF (www.gbif.org), iNaturalist (www.inaturalist.org), and BISON (www.bison.usgs.gov) are supported in ‘cRacle’ through the functions *gbif_get(), get_bison()*, and *inat()*. Each of these functions accepts arguments for a single taxon (genus, species, or family) to query and a maximum number of records to return. Data pre-processing, cleaning for outliers, spatial bias reduction, and climate data extraction for a set of georeferenced occurrence data is done through the *extraction()* function. Data downloading for multiple taxa and preprocessing functionality can be done in one step using the *getextr()* function, which is implemented with parallel computing capability to reduce user wait time.

### Likelihood Modeling

The primary modeling that CRACLE requires is the calculation of probability density functions using both parametric (Gaussian Normal) and non-parametric (Kernel Density) estimates. These calculations can be performed with ‘cRacle’ directly from the output of *extraction()* using the function(s) *dens_obj()* and *densform()* for multiple and single taxa respectively. Note that *dens_obj()* is simply a wrapper function for *densform()* designed to make multi-species likelihood model building simpler. Both parametric and non-parametric likelihood functions are always calculated and stored in the object produced by *densform()* and *dens_obj()*. This way all calculations are done in tandem for a dataset and results for both methods can be examined at the end of the CRACLE process.

The likelihood modeling carried out in the *densform()* function includes several options that can be manipulated by the user. The Kernel Density Estimation (KDE) employed by CRACLE can now be implemented with several kernel estimators including the standard Gaussian, Epanechnikov, cosine, optcosine, biweight, rectangular, and triangular kernels based on the R ‘stats’ package function *density()* (R Core Team, 2018). The kernel bandwidth can be optimized using standard criteria via the argument ‘bw’ in the *densform()* and *dens_obj()* functions. Likelihoods can be calculated based on a standard or conditional probability via the ‘manip’ argument where conditional probabilities take into account the background distribution of each climate parameter using a set of randomly sampled points from within a normalized distance from each primary occurrence location or from a user defined set of climate data. Finally, the resulting likelihood distributions can be trimmed to empirical ranges or confidence intervals to avoid unintended extrapolation using the ‘clip’ argument.

### CRACLE Joint Likelihood

After likelihood functions are estimated the next key CRACLE step is to calculate first the joint likelihood and then the maximum of the joint likelihood distribution. ‘cRacle’ implements the function *and_fun()* for calculating the joint likelihood and *or_fun()* for creating unions between taxa for merging likelihood features between groups (i.e., between species in a clade). The output of *and_fun()* is a set of joint likelihood features for all parameters, and this can be summarized by the function *get_optim()* which finds the optimal climate values for each parameter. The output of *get_optim()* provides the user with CRACLE results.

### Example Pseudocode

The general CRACLE modeling flow will use the following functions in order: *getextr() -> dens_obj()-> and_fun()-> get_optim();* to produce climate estimates from a list of taxa. This code would: download data and extract climate data, estimate likelihood distributions, calculate the joint likelihood, and find the optimum of the joint likelihood for each climate variable analyzed.

### Visualizing Likelihood and Joint Likelihood Functions

‘cRacle’ provides a set of functions for visualizing the likelihood functions as standard probability density distributions. These functions are *densplot()* for plotting single distribution objects (i.e., for one taxon/climate parameter pair), *addplot()* for adding to an existing *densplot()* figure, and *multiplot()* for plotting distributions for multiple taxa given a single climate parameter. For example, a user may plot the distributions for all taxa in their study for Mean Annual Temperature using *multiplot()* and the output from *dens_obj()*, and then add the joint likelihood function using *addplot()* and the model object output from *and_fun()*.

### Issue Tracking and Versioning

The repository at https://github.com/rsh249/cRacle.git will maintain the current development version of ‘cRacle’. Future major releases will be submitted to CRAN. Issues with running ‘cRacle’ code are accepted on the Git repository’s ‘Issues’ feature at https://github.com/rsh249/cRacle/issues or by email with the corresponding author of this publication.

### Data access

Primary biodiversity data, occurrence coordinates for your target taxa, and climate data are required for CRACLE modeling. We recommend primary biodiversity data downloaded from the Global Biodiversity Information Facility (GBIF) and support data access through ‘cRacle’ functions. Climate data must be in the form of a raster object readable by the R ‘raster’ package (Hijmans, 2018). Climate data from the WorldClim project (Hijmans et al., 2005; Fick and Hijmans, 2017) are recommended, but users may also use raster data downloaded from other sources such as CHELSA (Karger et al., 2017), and ENVIREM (Title and Bemmels, 2018).

### Estimation Climate from Vegetation – (ECV)

As originally described CRACLE consists of two methods for likelihood estimation: parametric and non-parametric probability functions. ‘cRacle’ implements these two approaches in tandem for each step of the likelihood analysis, and has implemented options about whether to calculate the likelihoods from standard probability or conditional probability. For comparison we have implemented calculations for the Mutual Climatic Range (MCR) method for ECV described by Thompson *et al.* (2012), though the unweighted MCR described is analogous to the Coexistence Approach (CA) described elsewhere (Mosbrugger and Utescher, 1997)

CRACLE (Harbert and Nixon, 2015) defines both a parametric and non-parametric probability density function for each taxon – climate parameter pair. CRACLE estimates likelihood for a set of taxa (t_1:i_). The CRACLE Parametric –Normal – probabilities are estimated as:

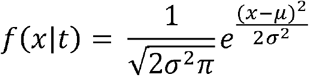

Whereas the CRACLE Non-Parametric – Kernel Density Estimation (KDE) – probabilities are estimated as:

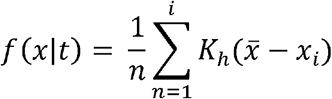

Where K is a Kernel function of properties XXXX, and ‘h’ is the kernel bandwidth, a smoothing parameter with value > 0. We recommend using either the “optcosine” or “gaussian” kernels and a bandwidth calculated using Silverman’s Rule of Thumb (Silverman, 1986). ‘cRacle’ allows for the calculation of probability density functions conditioned by the probability density of a random background sample for each climate parameter defined as:

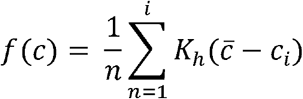

Where ‘c’ is the set of background climate given by a sample of points from within the study area. The conditional probability for a taxon (t) is defined as:

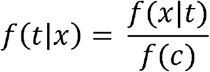

For the estimation of climate from vegetation CRACLE modeling calculates the joint likelihood for multiple taxon probability functions as:

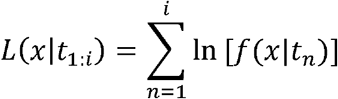

Where the maximum value of L(x| t_1:i_)corresponds to the value of the climate parameter ‘x’ that is most likely to lead to the coexistence of that set of taxa (t_1:i_).

Note that ‘cRacle’ also implements the calculation of a weighted mean by variance that can be substituted for the parametric CRACLE functions. Weighted means 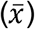 is calculated given a set of taxon means (x_1:i_) and standard deviations (σ_1:i_):

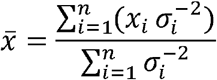

### Modern Validation

To test the performance of CRACLE that can be expected for paleoclimate modeling we set two experimental analyses to test aspects of the CRACLE modeling process.

First, we sample iNaturalist data (www.inaturalist.org) by a grid search of North America for plant species coexisting in 0.1×0.1 degree bounded search areas using the ‘rinat’ R package (Barve and Hart, 2017). Then we build CRACLE estimates for Mean Annual Temperature using the WorldClim 2.0 dataset at 2.5 arcminutes resolution (Fick and Hijmans, 2017) for parametric and non-parametric CRACLE as well as MCR and MCR un (unweighted) methods. Primary biodiversity data were obtained from GBIF (www.gbif.org) using the ‘cRacle’ *get_dist_all()* function and the GBIF RESTful JSON based API. Model overfitting for the iNaturalist dataset was tested by spatially thinning the raw GBIF data using the ‘cRacle’ *extraction()* function for factors of 2,4,6,8, and 10 times the dimensions of the 2.5 arcminute climate raster cells. The ‘cRacle’ thinning procedure is an imperfect, but efficient, spatial thinning method that is consistent with best-practices in Ecological Niche Modeling literature (Aiello-Lammens *et al.* 2015). CRACLE and MCR model output for each thinning level were summarized and reported for general guidance. The R code for the iNaturalist grid search test is available at https://github.com/rsh249/cracle_examples/inat_grid_search.

Second, the Botanical Information and Ecology Network (BIEN 4.0) database was queried for published vegetation plot surveys from North and South America using the ‘rbien’ R package (Maitner *et al.* 2017). For plot surveys with >5 plant taxa with distribution data available from GBIF we estimated climate parameters for all 19 bioclimatic variables from the WorldClim 2.0 data (Fick and Hijmans, 2017) using parametric CRACLE methods and spatial thinning to a factor of 2 times the 2.5 arcminute climate grid. Error in CRACLE are non-random (Harbert and Nixon, 2015), therefore non-linear regression modeling may be able to account for and correct common CRACLE errors. CRACLE results for 70,391 plot surveys were then partitioned 50:50 into training and testing sets by random sampling with replacement and Generalized Boosted Regression (GBR) models were fit to the training data for each of the 19 climate parameters using the R ‘gbm’ package (Greenwell *et al.* 2019). The GBR models made adjusted climate estimates using the CRACLE results in the test set to determine whether error patterns in CRACLE can be predicted by GBR and therefore may be corrected for in independent analyses. The R code for the iNaturalist grid search is available: https://github.com/rsh249/cracle_examples/cracle_bien.

## RESULTS

### Modern Validation

#### iNaturalist

The grid search of iNaturalist in North America yielded climate estimations by CRACLE (both parametric and non-parametric) and the Mutual Climatic Range (MCR) unweighted and weighted approaches for 285 sites at spatial thinning factors of 0, 2, 4, 6, 8, and 10 times the 2.5 arcminute climate raster. A total of 1,204,925 unique records were accessed from GBIF for this analysis for a total of 1541 species. The sampled localities ranged in mean annual temperature (MAT) from 0.4-26.1°C. The top performing (lowest error rates) estimates were the CRACLE non-parametric (‘kde’ in Figure 1) and the MCR weighted (‘mcr.w’ in Figure 1) methods. Mean errors for CRACLE were ~1°C and median errors were generally less than the means. Spatial thinning did not change mean or median error rates for any of the tested methods (Fig. 1B,C).

**Figure 1.**
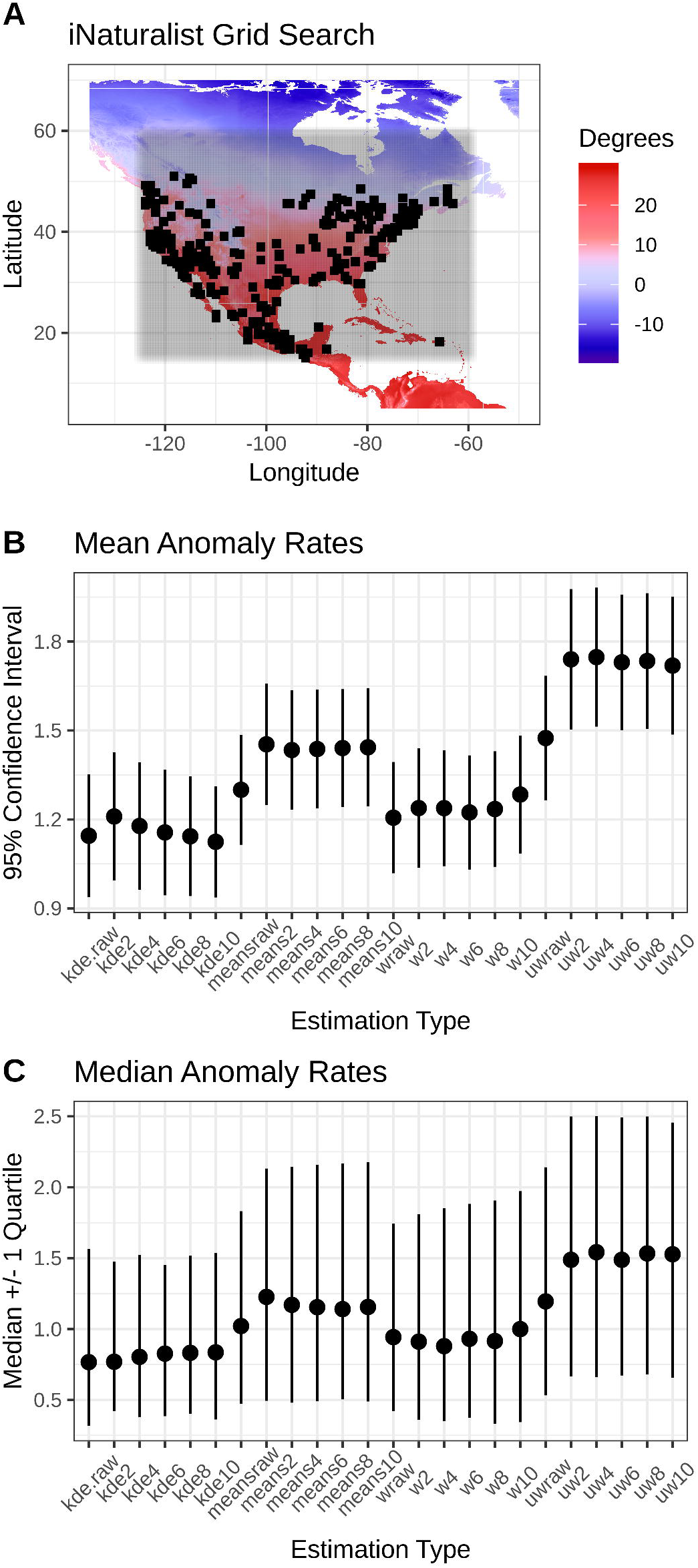
iNaturalist Grid Search Climate Estimation Results. A) Geographic search area (shading) with successful climate estimation locations marked (black squares). B) Mean Anomaly Rates, and C) Median Anomaly Rates for CRACLE (‘kde’ non-parametric, and ‘means’ parametric) and MCR (‘uw’ unweighted and ‘w’ weighted) results for factor levels of 0 (‘raw’), 2, 4, 6, 8, and 10 times the 2.5 arcminute climate raster.

#### BIEN

Non-parametric CRACLE estimates of all 19 WorldClim Bioclim variables (www.worldclim.org/bioclim) from the WorldClim 2.0 dataset (Fick and Hijmans, 2017) were generated for 70,391 vegetation plot surveys accessed through the BIEN 4.0 database using a total of 2,560,261 georeferenced specimen records from GBIF.

Mean absolute anomaly rates for temperature parameters were between 1-2°C with Pearson’s Correlation coefficients greater than 0.9 in the best scenarios (Table 1), but up to 5°C and correlation coefficients <0.7 for estimates of mean temperature during the wettest and driest quarters of the year (BIO8 and BIO9). The mean absolute anomaly rates for mean annual precipitation (MAP – BIO12) was 179mm with a correlation of 0.79 (Table 1).

**Table 1:** CRACLE performance statistics for 19 Bioclim parameters estimated for BIEN vegetation plot data (n=35,195 plots)

| Climate_Parameter | Mean Anomaly | Mean Absolute Anomaly | Median Absolute Anomaly | RMSE | NRMSE | Pearson's $r$ | Spearman's $r$ | Units |
| --- | --- | --- | --- | --- | --- | --- | --- | --- |
| wc2.0_bio_2.5m_01 | 0.34 | 1.40 | 1.09 | 1.86 | 0.05 | 0.95 | 0.96 | °C |
| wc2.0_bio_2.5m_02 | -0.31 | 1.01 | 0.74 | 1.39 | 0.07 | 0.82 | 0.79 | °C |
| wc2.0_bio_2.5m_03 | -0.25 | 2.51 | 1.80 | 3.44 | 0.04 | 0.87 | 0.87 | °C |
| wc2.0_bio_2.5m_04 | -23.93 | 64.64 | 42.35 | 92.68 | 0.06 | 0.84 | 0.85 | °C |
| wc2.0_bio_2.5m_05 | -0.19 | 1.71 | 1.19 | 2.32 | 0.07 | 0.92 | 0.95 | °C |
| wc2.0_bio_2.5m_06 | 0.64 | 2.13 | 1.57 | 2.93 | 0.05 | 0.91 | 0.92 | °C |
| wc2.0_bio_2.5m_07 | -0.94 | 2.42 | 1.97 | 3.19 | 0.07 | 0.79 | 0.83 | °C |
| wc2.0_bio_2.5m_08 | 1.00 | 5.60 | 3.27 | 7.81 | 0.18 | 0.69 | 0.63 | °C |
| wc2.0_bio_2.5m_09 | -1.00 | 5.64 | 2.43 | 8.80 | 0.16 | 0.60 | 0.63 | °C |
| wc2.0_bio_2.5m_10 | 0.03 | 1.30 | 0.97 | 1.74 | 0.06 | 0.94 | 0.96 | °C |
| wc2.0_bio_2.5m_11 | 0.31 | 1.89 | 1.36 | 2.66 | 0.05 | 0.92 | 0.93 | °C |
| wc2.0_bio_2.5m_12 | 26.20 | 179.17 | 111.69 | 275.09 | 0.06 | 0.79 | 0.83 | mm/year |
| wc2.0_bio_2.5m_13 | 7.32 | 25.04 | 11.31 | 47.92 | 0.07 | 0.70 | 0.75 | mm/month |
| wc2.0_bio_2.5m_14 | 9.89 | 15.49 | 9.72 | 21.79 | 0.10 | 0.77 | 0.76 | mm/month |
| wc2.0_bio_2.5m_15 | -7.11 | 11.66 | 8.30 | 16.12 | 0.15 | 0.69 | 0.71 | -- |
| wc2.0_bio_2.5m_16 | 21.84 | 68.96 | 31.48 | 135.63 | 0.08 | 0.69 | 0.76 | mm/3*month |
| wc2.0_bio_2.5m_17 | 26.80 | 45.61 | 27.49 | 64.92 | 0.08 | 0.80 | 0.76 | mm/3*month |
| wc2.0_bio_2.5m_18 | -6.20 | 34.76 | 23.16 | 52.69 | 0.05 | 0.90 | 0.91 | mm/3*month |
| wc2.0_bio_2.5m_19 | 34.97 | 99.35 | 73.22 | 142.96 | 0.10 | 0.57 | 0.69 | mm/3*month |

Generalized Boosted Regression (GBR) models were used to correct CRACLE estimates using independent random samples without replacement of 50% of the data for training and testing datasets. GBR correction improved CRACLE performance in the test set by 20-50% (Table 1, 2; Figure 2). CRACLE+GBM corrected anomalies are smaller and more symmetrically distributed around ‘0’ than the non-corrected CRACLE results (Figure 2D), and CRACLE+GBM estimates are more strongly correlated with the WorldClim data than the raw CRACLE results (Table 1, 2).

**Table 2:** GBR Corrected CRACLE performance statistics for 19 Bioclim parameters estimated for BIEN vegetation plot data (n=35,195 plots)

| Climate_Parameter | Mean Anomaly | Mean Absolute Anomaly | Median Absolute Anomaly | RMSE | NRMSE | Pearson's $r$ | Spearman's $r$ | Units |
| --- | --- | --- | --- | --- | --- | --- | --- | --- |
| wc2.0_bio_2.5m_01 | -0.06 | 1.11 | 0.85 | 1.51 | 0.04 | 0.96 | 0.96 | °C |
| wc2.0_bio_2.5m_02 | 0.00 | 0.84 | 0.60 | 1.18 | 0.06 | 0.86 | 0.81 | °C |
| wc2.0_bio_2.5m_03 | -0.01 | 2.08 | 1.48 | 2.89 | 0.04 | 0.91 | 0.89 | °C |
| wc2.0_bio_2.5m_04 | 1.03 | 51.65 | 36.94 | 72.73 | 0.05 | 0.90 | 0.87 | °C*100 |
| wc2.0_bio_2.5m_05 | -0.11 | 1.18 | 0.83 | 1.64 | 0.05 | 0.96 | 0.95 | °C |
| wc2.0_bio_2.5m_06 | -0.16 | 1.75 | 1.18 | 2.49 | 0.04 | 0.93 | 0.93 | °C |
| wc2.0_bio_2.5m_07 | -0.03 | 1.85 | 1.33 | 2.57 | 0.06 | 0.85 | 0.86 | °C |
| wc2.0_bio_2.5m_08 | -0.38 | 4.43 | 3.00 | 6.17 | 0.15 | 0.74 | 0.70 | °C |
| wc2.0_bio_2.5m_09 | 0.41 | 4.86 | 2.38 | 7.66 | 0.14 | 0.71 | 0.69 | °C |
| wc2.0_bio_2.5m_10 | -0.06 | 0.97 | 0.73 | 1.32 | 0.05 | 0.97 | 0.96 | °C |
| wc2.0_bio_2.5m_11 | -0.15 | 1.61 | 1.14 | 2.32 | 0.04 | 0.94 | 0.94 | °C |
| wc2.0_bio_2.5m_12 | 12.84 | 147.83 | 95.59 | 225.75 | 0.05 | 0.86 | 0.86 | mm/year |
| wc2.0_bio_2.5m_13 | 1.52 | 19.52 | 10.52 | 35.44 | 0.05 | 0.78 | 0.82 | mm/month |
| wc2.0_bio_2.5m_14 | 0.04 | 10.83 | 7.50 | 15.50 | 0.08 | 0.85 | 0.86 | mm/month |
| wc2.0_bio_2.5m_15 | 2.11 | 7.56 | 4.77 | 11.17 | 0.11 | 0.75 | 0.73 | -- |
| wc2.0_bio_2.5m_16 | 3.25 | 51.70 | 28.72 | 97.37 | 0.06 | 0.79 | 0.83 | mm/3*month |
| wc2.0_bio_2.5m_17 | -0.80 | 32.85 | 22.45 | 46.69 | 0.07 | 0.87 | 0.87 | mm/3*month |
| wc2.0_bio_2.5m_18 | 2.48 | 28.98 | 21.01 | 42.83 | 0.04 | 0.93 | 0.92 | mm/3*month |
| wc2.0_bio_2.5m_19 | 11.46 | 70.76 | 46.68 | 114.25 | 0.08 | 0.71 | 0.76 | mm/3*month |

**Figure 2.**
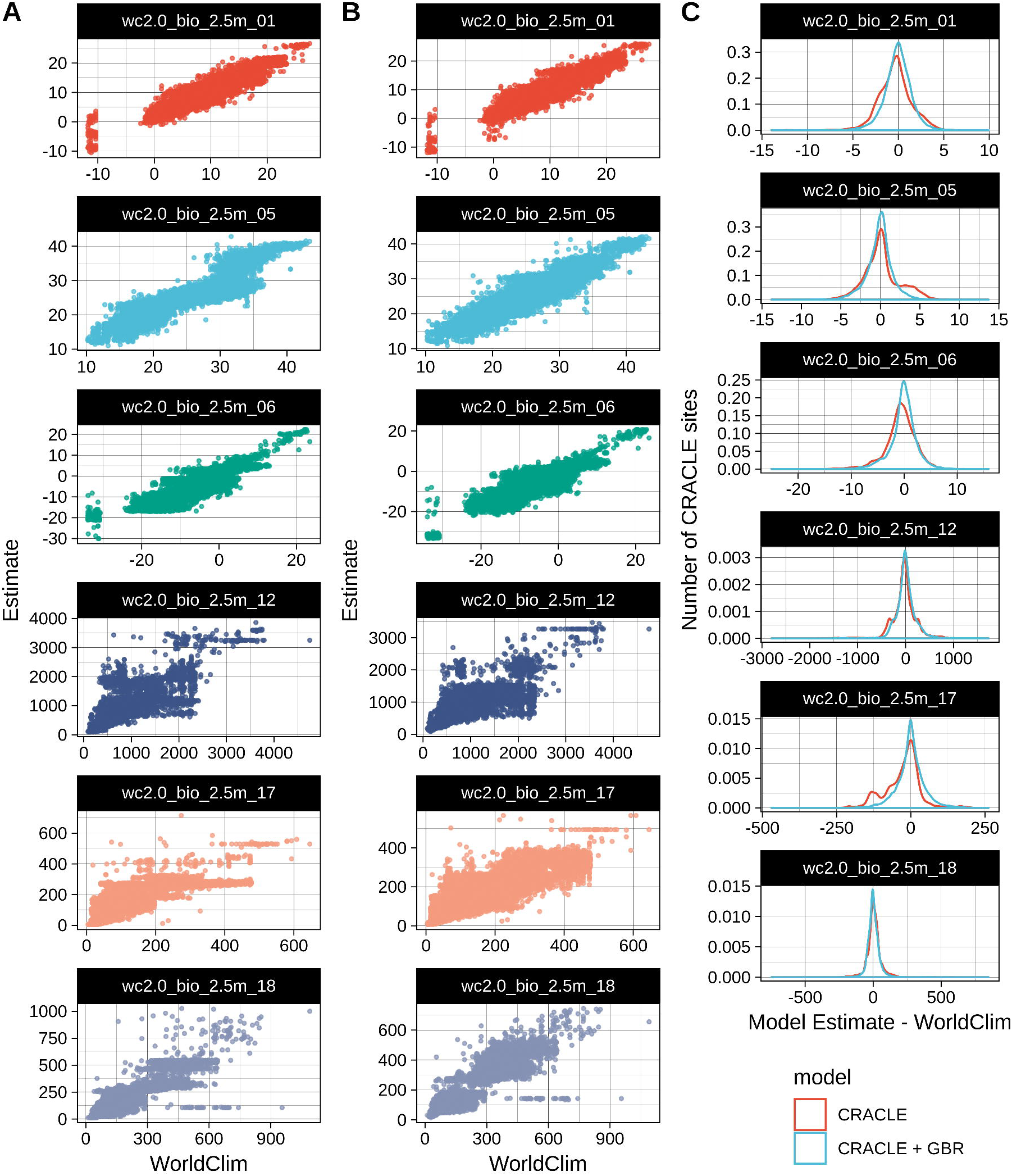
CRACLE estimates and GBR error correction demonstration for BIEN vegetation plot surveys and representative climate parameters. BIEN vegetation plot surveys from 70,391 unique localities (A) were analyzed with CRACLE (B) to estimate the 19 Bioclim variables (showing BIO1, 5, 6, 12, 17, and 18 here as top performing examples) from the WorldClim2.0 dataset (Fick and Hijmans, 2017). Generalized Boosted Regression error correction modeling was trained and tested (C) on independent subsets of plot data. GBR correction yielded overall reduced error rates and less biased estimates in many cases (D).

## DISCUSSION

The modern validation analyses conducted here should provide a good baseline for expected CRACLE performance going forward. More thorough testing in future studies is certainly welcome, but most modeling choices are should be made by the user to reflect any unique properties of their study system. The results we provide here indicate that the best estimates that are generated from the ‘cRacle’ software are the CRACLE non-parametric estimates with spatially thinned data, these yielding mean errors of approximately 0.5°C less than the Mutual Climatic Range (MCR) method, an analogous method to the widely used Coexistence Approach (Mosbrugger and Utescher, 1997). Furthermore, CRACLE results presented here compare favorably, though indirectly, to recent applications of the Weighted Average – Partial Least Squares (WA-PLS) modeling that is common for analysis of fossil pollen samples that uses modern analog communities to build proxy models (Montade *et al.*, 2019). A direct comparison of CRACLE and WA-PLS methods is necessary future work.

Through our analysis of BIEN hosted vegetation plot surveys we show that CRACLE can produce accurate climate estimates for a variety of both temperature and precipitation parameters (Figure 2), though some parameters are better predicted than others (Table 1). Notably, BIO8 (Mean Temperature of the Warmest Quarter), BIO9 (Mean Temperature of the Coldest Quarter), BIO13 (Precipitation of the Wettest Month), and BIO15 (Precipitation Seasonality – Coeff. Var.) are relatively poorly predicted by CRACLE, likely due to less direct impact of those variables on plant distributions. Whereas estimates of BIO1 (MAT), BIO5 (Max Temperature), BIO6 (Min Temperature), and BIO18 (Precipitation of the Driest Quarter) yield the highest correlation with known values (Table 1). We also show that CRACLE model correction via Generalized Boosted Regression (GBR) can lower error rates from the CRACLE baseline (Figure 2; Table 2). We expect that further regression model fitting on expanded datasets may yield further refinement, however, the GBR model correction is implemented in ‘cRacle’ *get_optim()* as an option for users to correct non-parameteric CRACLE estimates for the WorldClim 2.0 Bioclim variables only.

### Resources and Tutorials

We are actively developing resources and tutorials in support of the ‘cRacle’ R package. Demonstration code and short projects will be maintained on a continuing basis at https://github.com/rsh249/cracle_examples. We aim to provide a series of web tutorials for various CRACLE modeling tasks here to guide beginner users through the process. Tutorials, and issue tracking are distributed through the main ‘cRacle’ repository: https://github.com/rsh249/cRacle.git.

### Conclusion

‘cRacle’ is a new and actively maintained resource for climate estimation from biological community composition. These estimates are particularly relevant to the study of fossil systems where often the best indication of past climate is the community of plant and animal fossils. We show that users of ‘cRacle’ should expect estimates that are accurate (e.g., within 1°C for MAT) when applying best-practices.

## Acknowledgements

The authors would like to thank Stonehill College for their support of undergraduate research that made this study possible.

## AUTHOR CONTRIBUTIONS

AAB conceived of and implemented the iNaturalist grid search and CRACLE analysis and provided valuable testing, benchmarking, and feedback data on the ‘cRacle’ R library. RSH designed the BIEN vegetation plots CRACLE study and implemented the Generalized Boosted Regression model correction analysis. RSH designed and wrote the ‘cRacle’ R package code.

